# Foxp1 controls neural stem cell competence and bias towards deep layer cortical fates

**DOI:** 10.1101/386276

**Authors:** Caroline Alayne Pearson, Destaye M. Moore, Haley O. Tucker, Joseph D. Dekker, Hui Hu, Amaya Miquelajáuregui, Bennett G. Novitch

**Affiliations:** department of Neurobiology, David Geffen School of Medicine at UCLA, David Geffen School of Medicine at UCLA, Los Angeles, CA 90095, USA; Eli and Edythe Broad Center of Regenerative Medicine and Stem Cell Research, David Geffen School of Medicine at UCLA, Los Angeles, CA 90095, USA; Intellectual and Developmental Disabilities Research Center David Geffen School of Medicine at UCLA Los Angeles, CA 90095, USA; Molecular Biosciences University of Texas at Austin Austin, Texas 78712; Department of Microbiology School of Medicine University of Alabama at Birmingham Birmingham, AL 35205; Institute of Neurobiology University of Puerto Rico Medical Sciences Campus San Juan 00911, Puerto Rico; Lead Contact

**Keywords:** brain development, cerebral cortex, neurogenesis, neural stem cell, neural differentiation, Foxp1, transcriptional regulation

## Abstract

**SUMMARY:** The laminar architecture of the mammalian neocortex depends on the orderly generation of distinct neuronal subtypes by apical radial glia (aRG) during embryogenesis. We identify critical roles for Foxp1 in maintaining RG identity and gating the temporal competency for early neurogenesis. High levels of Foxp1 are associated with early aRG and are required to promote proliferation and influence cell division symmetry, favoring aRG expansion and production of early born neurons. The potent pro-progenitor functions of Foxp1 are revealed through its ability to preserve a population of cells with aRG identity throughout development and extend the early neurogenic period into postnatal life. Foxp1 further promotes the formation of cells resembling basal RG (bRG), a progenitor group implicated in the increased size and complexity of the human cortex. Consistent with this role, we show that FOXP1 is associated with the initial formation and expansion of bRG during human corticogenesis.

**HIGHLIGHTS:**

- Foxp1 is transiently expressed by aRG during the early phase of corticogenesis
- Foxp1 promotes self-renewing vertical cell divisions and aRG maintenance
- Foxp1 gates the time window of deep layer neurogenesis
- Ectopic Foxp1 expression can elicit bRG formation

## INTRODUCTION

The development of the central nervous system (CNS) depends on a pool of self-renewing neural stem cells being maintained over a protracted period of time to ensure the appropriate numbers of neurons, astrocytes, and oligodendrocytes are generated. Initially, the neural tube is composed of proliferative neuroepithelial progenitors that predominantly undergo symmetric and self-renewing divisions. In the developing neocortex, neuroepithelial progenitors transition into apical (ventricular) radial glia (aRG), which possess the ability to self-renew, directly generate neurons, or give rise to basal secondary progenitors, most notably short-lived intermediate progenitors (IP) and longer-lived basal radial glia (bRG) (Taverna et al., 2014). This transition marks the formation of two proliferative zones within the developing neocortex; the apical ventricular zone (VZ) composed of aRG and the adjacent subventricular zone (SVZ) where IP and bRG reside (Taverna et al., 2014). The increased size and complexity of the neocortex in higher mammals, particularly humans, is attributed to the enlargement of the SVZ resulting from expanded bRG formation (Lui et al., 2011; Florio and Huttner, 2014). Disruptions in neural progenitor maintenance and the balance between proliferative growth and differentiation are thought to underlie many neurodevelopmental disorders (Bae et al., 2015; Ernst, 2016); however, the genetic and cellular mechanisms governing these processes are not well understood.

The laminar organization of the mature neocortex is established during embryogenesis by the sequential generation of early-born deep layer neurons followed by later-born superficial layer neurons. A long-standing question has been the nature of the mechanisms that ensure that aRG produce specific types of neurons at appropriate times in development. Both viral and recombination-based lineage tracing studies have demonstrated that aRG present at the onset of neurogenesis have the capacity to give rise to all cortical neuron types (Gao et al., 2014; Ma et al., 2017). As neurogenesis proceeds, aRG lose their potential to give rise to deep layer neurons and begin to generate superficial layer neurons (Kwan et al., 2012; Greig et al., 2013; Dwyer et al., 2016). While this “temporal competency” model has had great appeal, it remains unclear how early aRG are biased to form deep layer neurons and what transition occurs to change their neurogenic output. Indeed, a contrasting model whereby deep vs. superficial layer neurons are produced by distinct progenitors rather than through temporal switching of a common pool has been proposed (Franco and Muller, 2013). In addition, it remains unknown how changes in progenitor output are coordinated with the mechanisms controlling progenitor maintenance.

Foxp proteins are a family of transcriptional repressors that are vital for the development of blood, lungs, heart, and the CNS (Wang et al., 2004; Hu et al., 2006; Shu et al., 2007; Dasen et al., 2008; Rousso et al., 2008; Rousso et al., 2012; Wang et al., 2014). Foxp1/2/4 are notably expressed throughout the developing cortex in both dividing progenitors and differentiated cell types (Ferland et al., 2003; Hisaoka et al., 2010; Araujo et al., 2015), and mutations have been linked to a range of cognitive disorders in humans including autism spectrum disorders (ASD) (Lai et al., 2001; Groszer et al., 2008; Hamdan et al., 2010; Horn et al., 2010; O’Roak et al., 2011; Le Fevre et al., 2013; Bacon et al., 2015; Sanders et al., 2015; Meerschaut et al., 2017). Previously, we have shown that the ability of Foxp4 to repress N-Cadherin is critical for directing the fate of newly differentiating neurons, regulating the detachment of motor neuron progenitors from the apical surface of the VZ in the spinal cord as well as formation of IP from aRG in the neocortex (Rousso et al., 2012). The function of Foxp proteins in dividing neural progenitors and their roles in temporal competency, however, has not been explored.

In this study, we demonstrate that Foxp1 levels influence the maintenance of aRG identity and the timing of neurogenic waves in the developing neocortex. Foxp1 is expressed at the outset of cortical neurogenesis and progressively declines as cells transition from deep to superficial layer neuron production. Gain or loss of Foxp1 function alters the plane of progenitor cell divisions and the balance between self-renewal and differentiation into secondary progenitors and neurons. Sustained Foxp1 expression can extend the formation of early born cell types into postnatal stages. We lastly show that Foxp1 is similarly expressed by aRG cells in the developing human brain, but in contrast to mouse, is also strikingly associated with bRG. Acute elevation of Foxp1 during the peak neurogenic period in the mouse cortex promotes the formation of ectopic bRG-like cells outside of the VZ, suggesting that Foxp1’s ability to promote RG identity may be utilized to generate and expand bRG in the human brain. Together, these studies define important roles for Foxp1 maintaining aRG identity, promoting the generation of deep layer neurons, and eliciting bRG formation.

## RESULTS

### Foxp1 is expressed by aRG and downregulated during the transition from deep to superficial layer neurogenesis

To assess the role of Foxp1 in cortical neurogenesis, we first analyzed its RNA expression during the peak period of neurogenesis in mice (E12.5-E15.5) to the onset of gliogenesis (E16.5) (Figures 1A and S1D). *Foxp1* is present at high levels in the VZ at E12.5 coincident with deep layer neurogenesis. At E13.5, *Foxp1* expression in the VZ becomes markedly reduced, whilst neurons in the cortical plate (CP) derived from the early progenitors continue to express high levels of *Foxpl*. The level of *Foxp1* in the VZ continues to decline as neurogenesis proceeds and is barely detectable by E16.5. When compared to other sites of *Foxp1* expression in the developing brain such as the lateral ganglionic eminences (LGE), the intensity of *Foxp1* in the VZ quantifiably diminishes from E12.5 to E15.5 and is not detectable above background at E16.5 (Figure S1D). Similar results were seen with Foxp1 antibody staining; Foxp1 protein is first detectable in VZ cells throughout the lateral cortex starting at E10.5, coinciding with the transition from neuroepithelial progenitors to aRG (Figure S1A). Foxp1 levels in the VZ plateau around E11.5- E12.5 and then markedly decline compared to other VZ-associated proteins such as Sox2, becoming undetectable by E16.5 (Figures 1B, S1B, and S1C). By contrast, Foxp1 protein levels in CP neurons appeared constant throughout the course of neurogenesis (Figure 1B).

**Figure 1.**
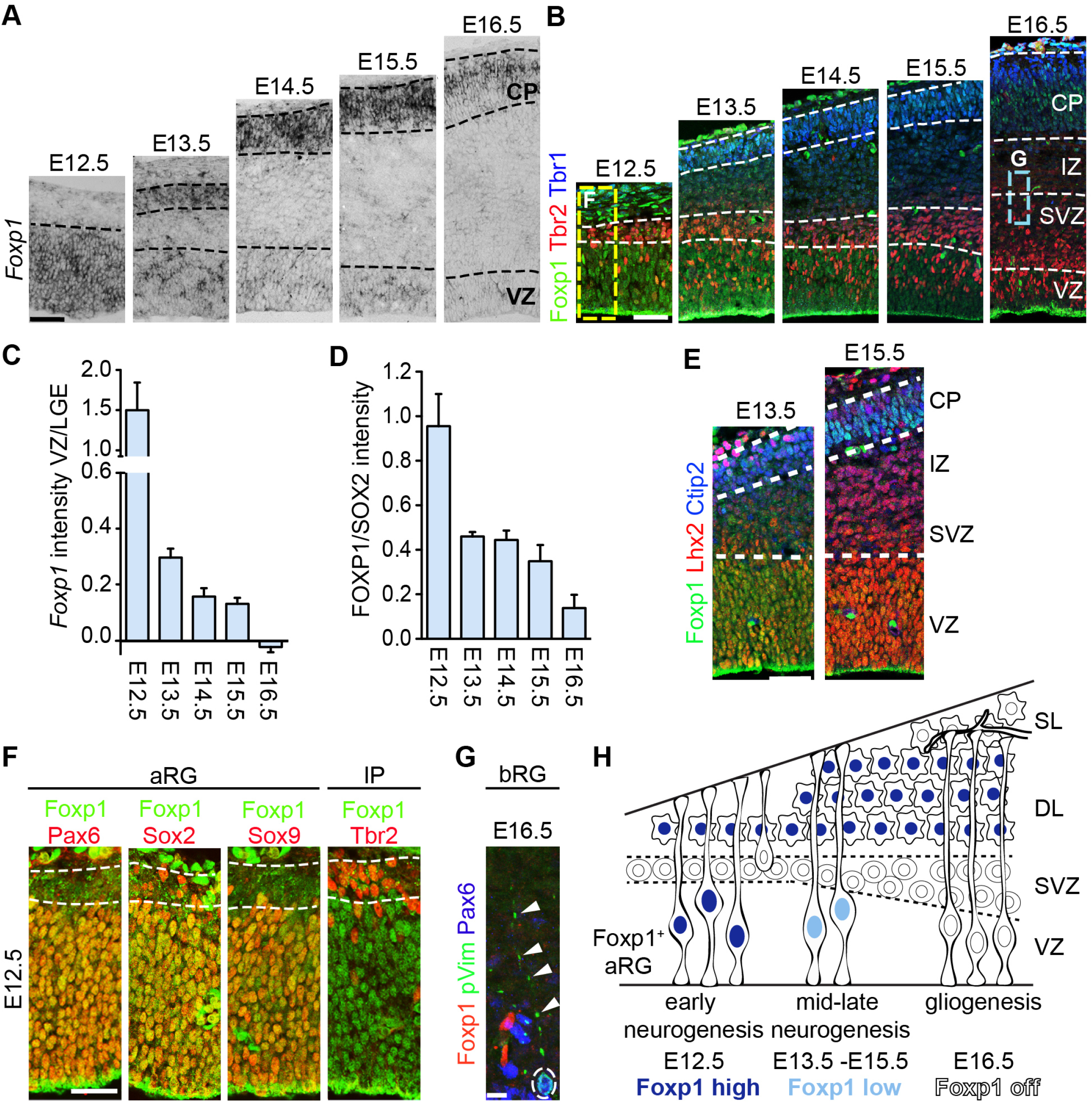
Foxp1 is expressed by aRG and downregulated during the transition from deep to superficial layer neurogenesis. (A)*Foxp1* mRNA expression in the VZ and CP during mouse cortical neurogenesis (E12.5-E16.5).(B) Immunostaining for Foxp1 protein in the VZ and CP. Tbr2 and Tbr1 immunostaining demarcate the SVZ and CP, respectively. Dashed boxes indicate areas of additional analysis displayed in panels F and G. (C) Plot of the relative expression of Foxp1 mRNA in the VZ compared to levels in the lateral ganglionic eminence (LGE; Figure S1D). Mean intensity values ± SEM were calculated from at least 3 sections per group from multiple embryos. (D) Quantification of Foxp1 protein levels in comparison to Sox2 levels (Figure S1B and S1C). Relative intensity ± SEM calculated from at least 3 sections per group from multiple embryos. Note Foxp1 staining along the apical edge of the VZ is non-specific (Figure S2C). (E) Immunostaining for Foxp1, Ctip2 and Lhx2 in the cortex at E13.5 and E15.5. Dashed white lines demarcate the approximate boundaries of the VZ and CP. (F) Immunostaining for Foxp1 showing overlap with the aRG markers Pax6, Sox2, Sox9, but not with the IP marker Tbr2. (G) Mouse bRG labeled with phosphorylated Vimentin and Pax6 do not express Foxp1. Arrowheads indicate basal process; white dashed circle denotes the mitotic cell soma. (H) Summary of Foxp1 expression in the VZ; high levels (dark blue) are expressed early during deep layer neurogenesis (E10.5-E12.5). Early born neurons also express Foxp1. As development proceeds (E13.5-E16.5), Foxp1 levels decrease in aRG (light blue) during superficial layer neurogenesis and are absent initiation of gliogenesis. High levels of Foxp1 are retained by deep layer neurons. aRG, apical radial glia; bRG, basal radial glia; VZ, ventricular zone; SVZ, subventricular zone; DL, deep layers; SL, superficial layers. Scale bars represent 50 μm in panels A, B and E, 25 μm in F, and 10 μm in G. See also Figure S1.

Within the VZ, Foxp1 expression overlaps with several well-established markers of aRG cells including Sox2, Sox9, and Pax6 (Figure 1D). Foxp1 is also expressed in early born, deep layer cortical neurons marked by Tbr1 expression, yet excluded from IP in the SVZ demarcated by Tbr2 expression (Figures 1A and 1D). High levels of Foxp1 within VZ progenitors notably concur with the peak period of deep layer production, marked by the appearance of Tbr1+ and Ctip2+ cells in the CP (Figures 1A and 1C). The observed decline in Foxp1 expression starting around E13.5 coincides with the production of Lhx2+ superficial layer neurons (Figure 1C). To identify bRG cells outside of the VZ, we costained for Pax6 and phosphorylated Vimentin (pVim) at a developmental stage that bRG are most abundant (E16.5; see dashed box in Figure 1B), yet found no evidence of Foxp1 in cells positive for these markers (Figure 1E). Collectively, these data show that Foxp1 expression is selectively expressed by aRG during the period of deep layer neurogenesis early in development, and progressively lost from these cells as they transition to superficial layer neurogenesis and, thereafter, gliogenesis (Figure 1F).

### Foxp1 promotes radial glial maintenance

To determine the roles that Foxp1 plays in aRG during cortical development, we conditionally elevated or ablated Foxp1 expression by crossing mice carrying *Emx1^Cre^* alleles, which drive recombination in neuroepithelial progenitors starting at ~E10, to those harboring either a Cre-inducible transgene driving Foxp1 expression from the CAG enhancer/promoter (“Foxp1^ON^” condition) or a Cre-inactivatable Foxp1 allele (“Foxp1^OFF^” condition) (Figure S2). *Emx1^Cre^*-negative littermate controls were used in all experiments. In E13.5 Foxp1^ON^ cortices, the number of Pax6+ aRG in the VZ increased whilst the number of Tbr2+ IP in the SVZ was reduced (Figures 2A, 2B, and 2F). Conversely, in Foxp1^OFF^ cortices there was a decrease in the number of aRG and corresponding increase in SVZ IP (Figures 2A, 2B, and 2F). On average, we observed a 2.5:1 ratio of Pax6+ aRG to Tbr2+ IP in control cortices. This ratio increased to 3.4:1 in Foxp1^ON^ cortices and decreased to less than 2:1 in Foxp1^OFF^ embryos (Figure 2G). These effects on progenitor identities coincided with changes in the production of Tbr1+ neurons at E13.5, with a reduction seen in Foxp1^ON^ cortices and increase in Foxp1^OFF^ cortices (Figures 2C and 2H). The production of the earliest born preplate neurons was similarly impacted by Foxp1 manipulation, with fewer Calretinin+ cells present in the mantle zone of Foxp1^ON^ cortices and more cells in Foxp1^OFF^ specimens (Figures S2E and S2F). Together, these data show that Foxp1 promotes the maintenance of aRG and inhibits both IP and neuron formation.

**Figure 2.**
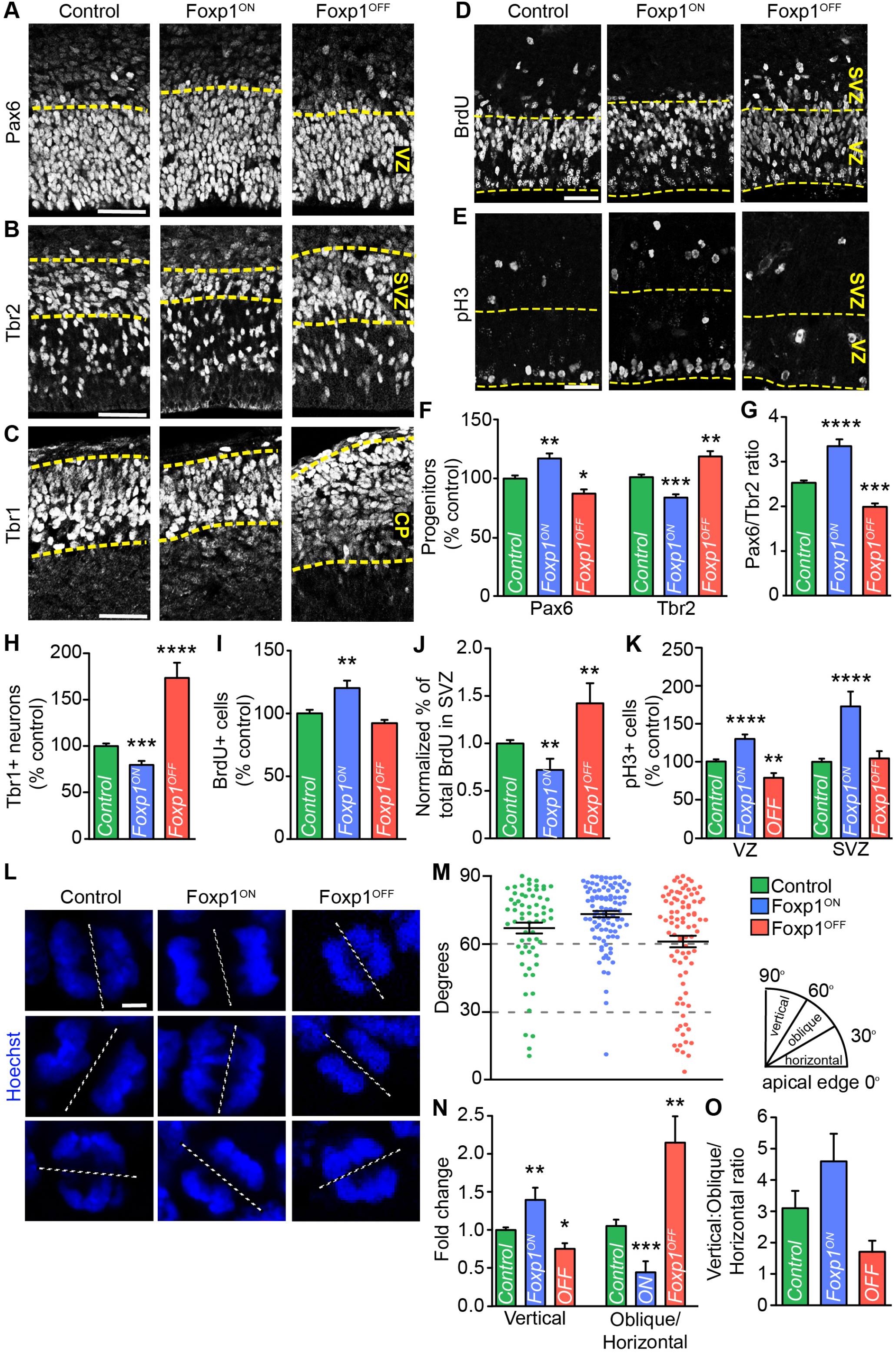
Foxp1 promotes radial glia identity. (A-C) Immunostaining for Pax6 in VZ aRG, Tbr2 in SVZ IP, and Tbr1 in CP neurons in E13.5 control, Foxp1^ON^ and Foxp1^OFF^ embryos. Yellow dashed lines indicate the approximate boundary of each compartment of the developing cortex. (D-E) BrdU incorporation (2-hour pulse) and immunostaining for phosphorylated Histone H3 (pH3) in E13.5 control, Foxp1^ON^ and Foxp1^OFF^ embryos. (F, H) Relative number of Pax6+ aRG, Tbr2+ IP, and of Tbr1+ CP neurons in control, Foxp1^ON^ and F oxp1^OFF^ embryos. (G) Quantification of the ratio of Pax6+ aRG to Tbr2+ IP. (I) Relative number of BrdU-labeled cells in control, Foxp1^ON^ and Foxp1^OFF^ embryos. (J) Percentage of BrdU+ cells located within the SVZ of control, Foxp1^ON^ and Foxp1^OFF^ embryos. (K) Relative number of phosphorylated Histone H3 (pH3+) cells. (L) Examples of the division angle of apical cells in anaphase detected by Hoechst staining. Dashed lines denote plane of division in relation to the apical surface. Vertical division 60-90°, oblique division 30-60°, horizontal division 0-30°. (M) Scatter plot analysis of division angle in control, Foxp1^ON^ and Foxp1^OFF^ conditions. Each dot represents a cell. Mean ± SEM from 80-100 cells analyzed from at least 5 embryos per condition. (N) Fold change differences in the mitotic behavior of dividing aRG upon Foxp1 manipulation. (O) Graphical representation of the vertical to oblique/horizontal division ratio in control, Foxp1^ON^ and F oxp1^OFF^ embryos at E13.5. Unless otherwise indicated, plots above display mean ± SEM from multiple sections taken from at least 3 embryos per condition. Significance was determined by Student’s t-test, *p <0.05; **p <0.005; ***p <0.001; ****p <0.0001. VZ, ventricular zone; SVZ, subventricular zone; CP, cortical plate. Scale bars represent 50 μm in panels A-E and 2 μm in L. See also Figure S2.

### Foxp1 promotes vertical, self-renewing cell divisions

Given its role in aRG maintenance, we next asked whether Foxp1 influences proliferation and selfrenewal. To determine whether Foxp1 promotes proliferation, we analyzed Bromodeoxyuridine (BrdU) incorporation in proliferating cells in our genetically modified mouse lines. In Foxp1^ON^ embryos, the total number of BrdU+ cells in the cortex increased whereas Foxp1^OFF^ embryos showed a small albeit not statistically significant decrease (Figures 2D and 2I). Whilst the total number of BrdU+ cells in Foxp1^OFF^ animals was not overtly affected, the distribution of BrdU+ cells in the VZ vs. SVZ was nevertheless altered (Figure 2D; VZ and SVZ demarcated by yellow dashed lines). We therefore analyzed the distribution of BrdU+ cells between the VZ and SVZ using Pax6 to demarcate the position of the VZ (Figure S3), and found that the percentage of BrdU+ cells in the SVZ decreased in Foxp1^ON^ embryos while it increased in Foxp1^OFF^ embryos (Figure 2J). These results suggest that aRG in Foxp1^ON^ cortices are more proliferative and thereby retained within the VZ. By contrast, aRG in Foxp1^OFF^ cortices are either rapidly differentiating and exiting the VZ immediately after taking up the BrdU label, or forming SVZ IP that are more proliferative than those in control animals. To discriminate between these two possibilities, we analyzed the number of mitotic phosphorylated Histone H3+ (pH3+) cells. In Foxp1^OFF^ cortices there were reductions in VZ mitoses but no significant changes in the SVZ (Figures 2E and 2K). Together with our analyses of IP and neuron numbers, these data suggest that in the absence of Foxp1, aRG have an increased propensity to exit the VZ and differentiate. By contrast, Foxp1 elevation leads to more VZ and SVZ mitoses (Figures 2E and K).

Several studies have suggested that during early phases of neurogenesis the angle and symmetry of neural progenitor cells divisions is tightly regulated such that the majority of cells undergo vertical (mitotic spindle angle of 60-90° relative to the apical plane), self-renewing or direct neurogenic divisions (i.e. producing one aRG and one neuron). However, as aRG transition to late neurogenesis this regulation becomes relaxed resulting in an increase in oblique/horizontal divisions (30-60°/0-30°) that favor the generation of basal progenitors and indirect neurogenesis (Lancaster and Knoblich, 2012). We accordingly measured mitotic spindle angles in Foxp1-manipulated cortices at E13.5, a time at which both modes of division can be readily seen (Figure 2L). In control cortices, the majority of aRG divisions appeared to be vertical (mean angle of 67°) with a minority fraction appearing horizontal, resulting in a 3:1 vertical to oblique/horizontal division ratio (Figure 2L, 2M, and 2O). In Foxp1^ON^ mice, mitoses were even more biased towards vertical divisions, with a mean division angle of 73° and 1.4-fold increase in the incidence of vertical divisions, raising the vertical to oblique/horizontal ratio to 4.5:1 (Figures 2M-2O). Foxp1^OFF^ cortices displayed the opposite trend, with a reduction in the mean angle of divisions to 62° and > 2-fold increase in the frequency of oblique/horizontal mitoses at the expense of vertical events (Figures 2M and 2N). The ratio of vertical to oblique/horizontal divisions was correspondingly reduced to 1.7:1 (Figure 2O).

Combined with our analysis of progenitor subtypes, these data show that Foxp1 promotes vertical, symmetric self-renewing aRG divisions that are characteristic of the early phase of cortical neurogenesis. Elevating Foxp1 biases cells towards vertical cell divisions which enlarges the aRG compartment and decreases both IP and neuron formation. In contrast, Foxp1 loss leads to more oblique aRG divisions, resulting in decreased aRG maintenance and precocious production of IP and neurons.

### Foxp1 manipulation impacts deep vs. superficial layer neurogenesis

As Foxp1 expression levels in aRG differ between the periods of deep vs superficial layer neurogenesis, we examined how Foxp1 misexpression or deletion from progenitors impacts the establishment of different cortical layers at E18.5 (schematic Figure 3A). In Foxp1^ON^ embryos, we found that Ctip2^high^ layer V neurons were significantly increased, as were Tbr1+ layer VI neurons (Figures 3C, 3D, and 3F). By contrast, Cux1+ and Lhx2+ layer II/III and IV neurons were reduced (Figures 3B, 3E, and data not shown). The ratio of deep to superficial layer neurons accordingly increased, from 2:1 in controls to 4.4:1 in Foxp1^ON^ cortices (Figure 3G). Early Foxp1 inactivation had the opposite effect, with deficits in both Ctip2^high^ layer V neurons and Tbr1+ layer VI neurons (Figures 3C, 3D, and 3F). However, the total number of Cux1+ and Lhx2+ layer II/III and IV did not appear to be affected (Figures 3B, 3E, and data not shown). Consequently, the ratio of deep to superficial layer neurons decreased to 1.7:1 in Foxp1^OFF^ cortices (Figure 3G).

**Figure 3.**
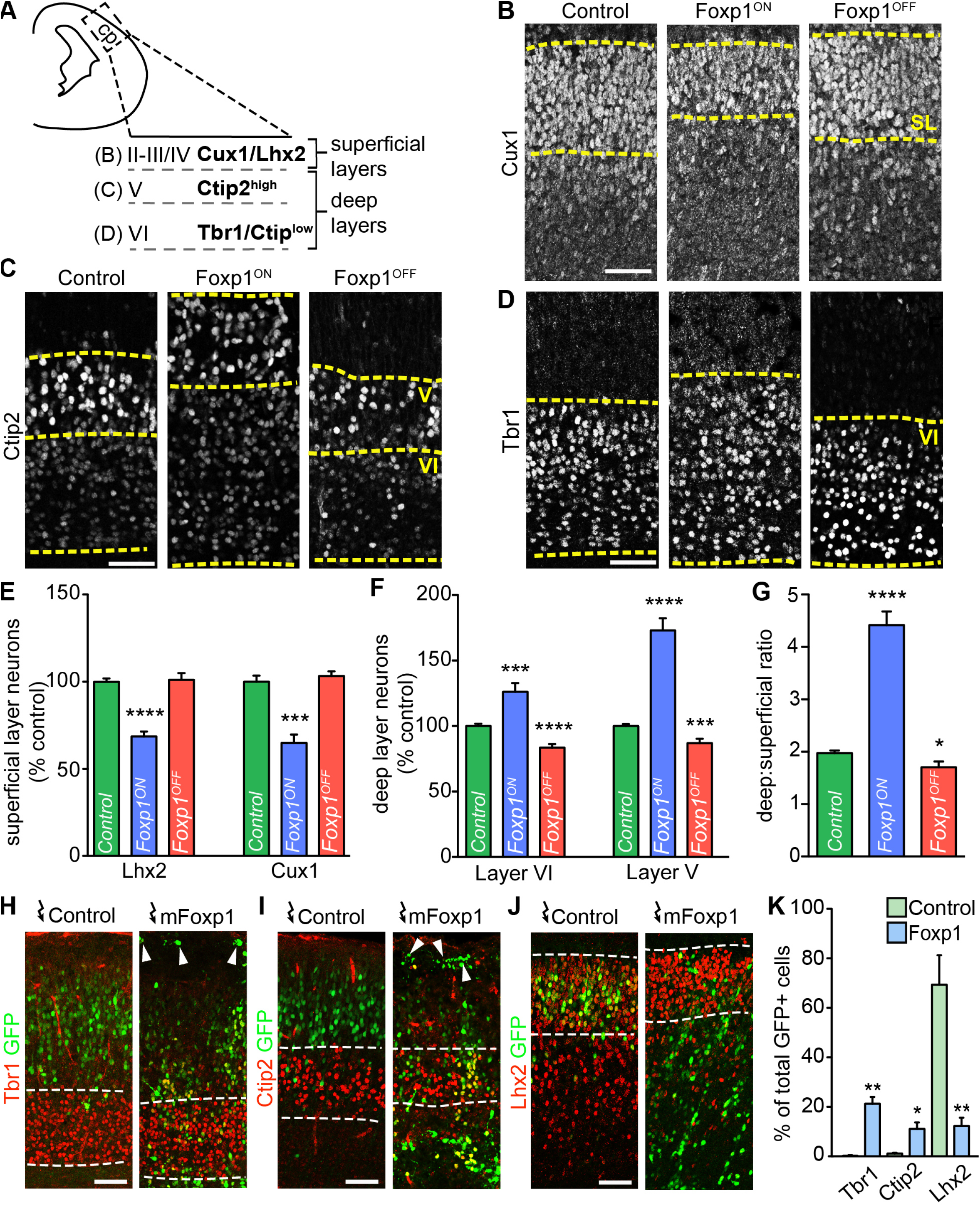
Foxp1 manipulation impacts deep vs. superficial layer neurogenesis. (A) Schematic of the lateral cortex area analyzed at E18.5. Cux1 and Lhx2 were used to delineate superficial layers, and Tbr1 and Ctip2 for deep layers. (B-D) Immunostaining for Cux1, Tbr1, and Ctip2 in control, Foxp1^ON^ and Foxp1^OFF^ cortices. Dashed yellow lines demarcate the approximate boundaries of the indicated layers. SL, superficial layers (II/III-IV). (E-F) Relative number of deep and superficial layer neurons in E18.5 control, Foxp1^ON^ and Foxp1^OFF^ conditions. Mean ± SEM from multiple sections from 3-6 embryos per condition are shown. (G) Graphical representation of the ratio of deep layer (VI/V; Tbr1+ Ctip2^high^) to superficial layer neurons. (H-J) Immunostaining for Tbr1, Ctip2, and Lhx2 in E18.5 cortices electroporated at E13.5 with IRES-GFP control or mouse Foxp1-IRES-GFP expression plasmids. (K) Quantification of the layer marker expression within the electroporated cells. Mean ± SEM from multiple sections from 2-3 animals per condition are shown. Scale bars represent 50 μm in all panels. Significance was determined using a Student’s t-test, *p <0.05; **p <0.005; ***p <0.001; ****p <0.0001. See also Figures S3 and S4.

In addition to these specific changes in neuronal fates, we found that the overall numbers of CP neurons, demarcated by expression of the pan-neuronal marker Mytl1, were slightly reduced in both Foxp1^ON^ and Foxp1^OFF^ embryos (Figures S3A and S3B). These deficits most likely reflect enhanced maintenance and reduced differentiation of aRG progenitors under the Foxp1^ON^ condition compared to precocious differentiation and depletion of the aRG pool in Foxp1^OFF^ cortices. Apoptosis, as evidenced by cleaved Caspase3 staining, did not appear to be grossly impacted by either Foxp1 manipulation (Figure S3C).

To confirm that the observed changes in neuronal identities and layer formation were consequences of upregulating Foxp1 in progenitors rather than neurons, we compared our findings based on *Emx1*^Cre^-mediated activation to those achieved using *Nex1^Cre^* mice, where Cre-mediated recombination is restricted to postmitotic neurons (Goebbels et al., 2006; Figure S3D). In *Nex1^Cre^;* Foxp1^ON^ cortices, we observed small reductions rather than gains in Ctip2^high^ layer V neurons, and no changes in the formation of either Tbr1+ layer VI or Cux1+ layer II/III-IV cells (Figures S3E-G) as was seen in Emx1^Cre^; Foxp1^ON^ animals (Figure 4). Thus, the influence of Foxp1 manipulation on neuronal fate specification appears due more to its actions in neural progenitors rather than differentiated neurons.

**Figure 4.**
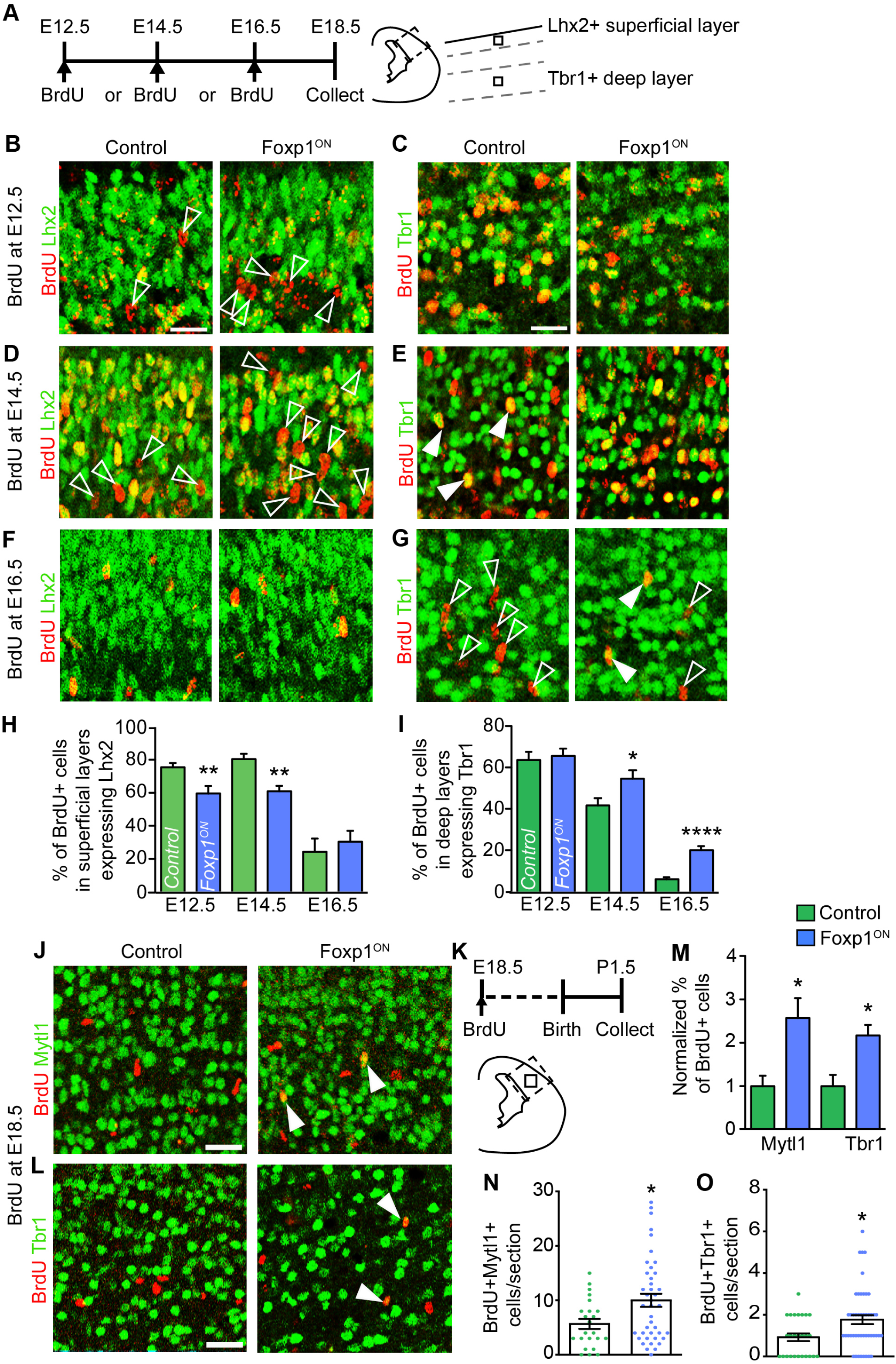
Sustained Foxp1 expression prolongs the period of deep layer neurogenesis. (A) Schematic of experimental design. Pregnant dams were injected with BrdU at E12.5, E14.5, or E16.5 and embryos collected at E18.5. Dashed box denotes area analyzed in subsequent panels. Tbr1 and Lhx2 were used to identify deep and superficial layers respectively. Small boxes represent areas enlarged in panels B-G. (B-G) Immunostaining for BrdU and layer markers in control versus Foxp1^ON^ embryos pulse labeled with BrdU at the indicated time points. Clear and filled arrowheads respectively denote cells staining for BrdU alone (red) or expressing the indicated layer markers (yellow). (H) Quantification of the percentage of BrdU-labeled cells expressing Lhx2 or Tbr1 within the superficial or deep layers of the cortex, respectively, after injection at E12.5, E14.5 or E16.5 time points. Plot displays mean ± SEM from multiple sections from at least 7 embryos per time point for each condition. (J-M) Late stage BrdU birthdating was accomplished by injection of BrdU into pregnant dams at E18.5 and collection of newborn pups at P1.5 (see experimental outline in panel K). BrdU-labeled cells expressing the pan neuronal marker Mytl1 or deep layer marker Tbr1 were scored and results for control and Foxp1^ON^ pups plotted in M. Plots display mean ± SEM from multiple sections from at least 4 pups per condition. Significance was determined using a Mann Whitney test, *p <0.05. (N-O) Scatter plot analysis of the number of BrdU+ Mytl1+ and BrdU+ Tbr1+ cells per section in control and Foxp1^ON^ conditions. Mean ± SEM from ~35 sections from at least 4 pups per condition is shown. Significance was determined using a Student’s t-test (H-I) or Mann-Whitney test (M-O). *p <0.05; **p <0.005; ***p <0.001; ****p <0.0001. Scale bars represent 20 μm in all panels. See also Figure S4.

Since Foxp1 manipulation altered the course of neurogenesis, we asked whether it also impacted gliogenesis by costaining for Brain lipid-binding protein (BLBP, also known as Fabp7) and Sox2 to identify astrocyte progenitors in E18.5 embryos. Only BLBP+ Sox2+ cells outside of the VZ/SVZ were counted to distinguish these cells from aRG which also express these markers. BLBP+ Sox2+ cells were significantly decreased in Foxp1^ON^ embryos, while there were no changes in Foxp1^OFF^ animals (Figures S5D-S5F and data not shown). Together, these data demonstrate that high levels of Foxp1 in aRG promote the generation of deep layer neurons at the expense of superficial layer neurons and glia, whilst the loss of Foxp1 leads to selective deficits in early but not late-born cells.

### Foxp1 elevation prolongs the period of deep layer neurogenesis

The observed changes to neuronal fate following *Foxp1* manipulation raised the question of whether sustained Foxp1 expression might extend the capacity of late aRG to give rise to deep layer neurons beyond their normal period of formation. To address this possibility, we examined the fate of cells in cortices electroporated with either control nEGFP or Foxp1-IRES-nEGFP expression constructs at E13.5, the stage at which *Foxp1* expression starts to diminish and aRG are transitioning to superficial layer neurogenesis. Pups were collected at birth (P0.5) and the distribution of GFP+ cells throughout the CP was analyzed using Tbr1 and Ctip2 to distinguish neurons in layers VI and V, and Lhx2 for cells in layers II/III-IV (Figure 3H). After control vector electroporations, GFP+ cells were predominantly localized in superficial layers (demarcated by condensed Lhx2+ cells; white dashed lines; Figures 3H and 3I), with >70% of GFP+ cells expressing Lhx2. By contrast, cells transfected with Foxp1-IRES-nEGFP plasmid were distributed throughout the CP (Figure 3H). There was also a significant increase in the percentage of GFP+ cells expressing Tbr1 (from 0.3% to 21%) and Ctip2 (from 1% to 11%), and a concomitant decrease in the percentage of GFP+ cells expressing Lhx2 (Figures 3H and 3I).

In cortices electroporated with Foxp1, GFP+ cells were frequently detected in the outermost layers, and many expressed Calretinin, a marker associated with the earliest born cells in layer I, a result that was not observed with the nEGFP control vector (arrowheads in Figures 3H and S5C). Many GFP+ neurons expressing Ctip2 and Tbr1 were also detected outside of their normal location within layers V and VI in Foxp1-electroporated cortices but not in control pups (Figures S5A and S5B; see controls in Figure 4H). These ectopic neurons were often clustered together in structures resembling heterotopias.

To further characterize Foxp1’s capacity to prolong the generation of early born deep layer neurons, we performed neuronal birthdating analysis by injecting pregnant Foxp1^ON^ dams with BrdU at E12.5, E14.5 or E16.5 and collecting embryos at E18.5. We subsequently costained cells with antibodies against BrdU and either Tbr1 or Lhx2 as indicators of early-born deep vs. later-born superficial neurons, respectively (Figure 4A). In control cortices, BrdU injections at E12.5 labeled approximately 60% of deep layer neurons born at that time point (Figures 4C and 4I). The frequency of BrdU labeling of deep layer neurons significantly dropped at E14.5, and were rare (< 5%) at E16.5 (Figures 4E, 4G, and 4I). As expected, superficial layer neurons were readily labeled by BrdU injections at either E12.5 or E14.5 with a steep decline seen at E16.5, reflecting the end stages of cortical neurogenesis (Figures 4B, 4D, 4F, and 4H). In Foxp1^ON^ embryos, we observed a marked increase in the birth of deep layer neurons labeled by BrdU injections at E14.5 or E16.5 (Figures 4C, 4E, 4G, and 4I). Conversely, the percentage of superficial layer neurons labeled by BrdU administration at E12.5 or E14.5 was reduced (Figures 4B, 4D, 4F, and 4H). The birth of BLBP+ astrocyte progenitors marked by BrdU labeling at E16.5 was similarly suppressed (Figures S5G and S5H).

We further asked whether Foxp1 maintenance could prolong deep layer neurogenesis into postnatal life by injecting pregnant dams with BrdU at E18.5 and collecting pups just after birth (P1.5; Figure 4K). Costaining analysis of BrdU with Mytl1 revealed that a small number of neurons are born during this period in control pups (average 5.7 ± 0.9 cells per section, maximum 15). This number increased to an average of 10 ± 1.2 cells per section, maximum 28 in Foxp1^ON^ pups, along with an overall increase in the percentage of BrdU+ cells that were Mytl1+ (Figures 4J, 4M and 4N). Analysis of BrdU+ cells within the deep layers further demonstrated that there was a >2-fold increase in both total numbers and percentage of BrdU+ Tbr1+ cells (Figures 4L, 4M, and 4O). Collectively, these data demonstrate that sustained Foxp1 expression can extend the neurogenic period and permit the generation of deep layer neurons into late embryogenesis and postnatal life, while impeding superficial layer neurogenesis and gliogenesis.

### Foxp1 can sustain an aRG-like population into adulthood

We next examined how Foxp1’s ability to promote aRG identity impacts other progenitor populations at later stages of development. In E16.5 Foxp1^ON^ cortices, there was a notable increase in the number of Pax6+ cells in the VZ and coincident loss of Tbr2+ IP in the SVZ (Figures 5A and 5B). These effects appeared to be cumulative; whereas we had found a 16% increase in Pax6+ numbers at E13.5, a 26% increase was seen at E16.5 along with a reciprocal decrease in the number of Tbr2+ IP (Figures 2F, 5A, and 5B). At this later stage, the ratio of Pax6+:Tbr2+ cells was 2:1 in Foxp1^ON^ cortices compared 1.5 in controls (Figure 5C).

**Figure 5.**
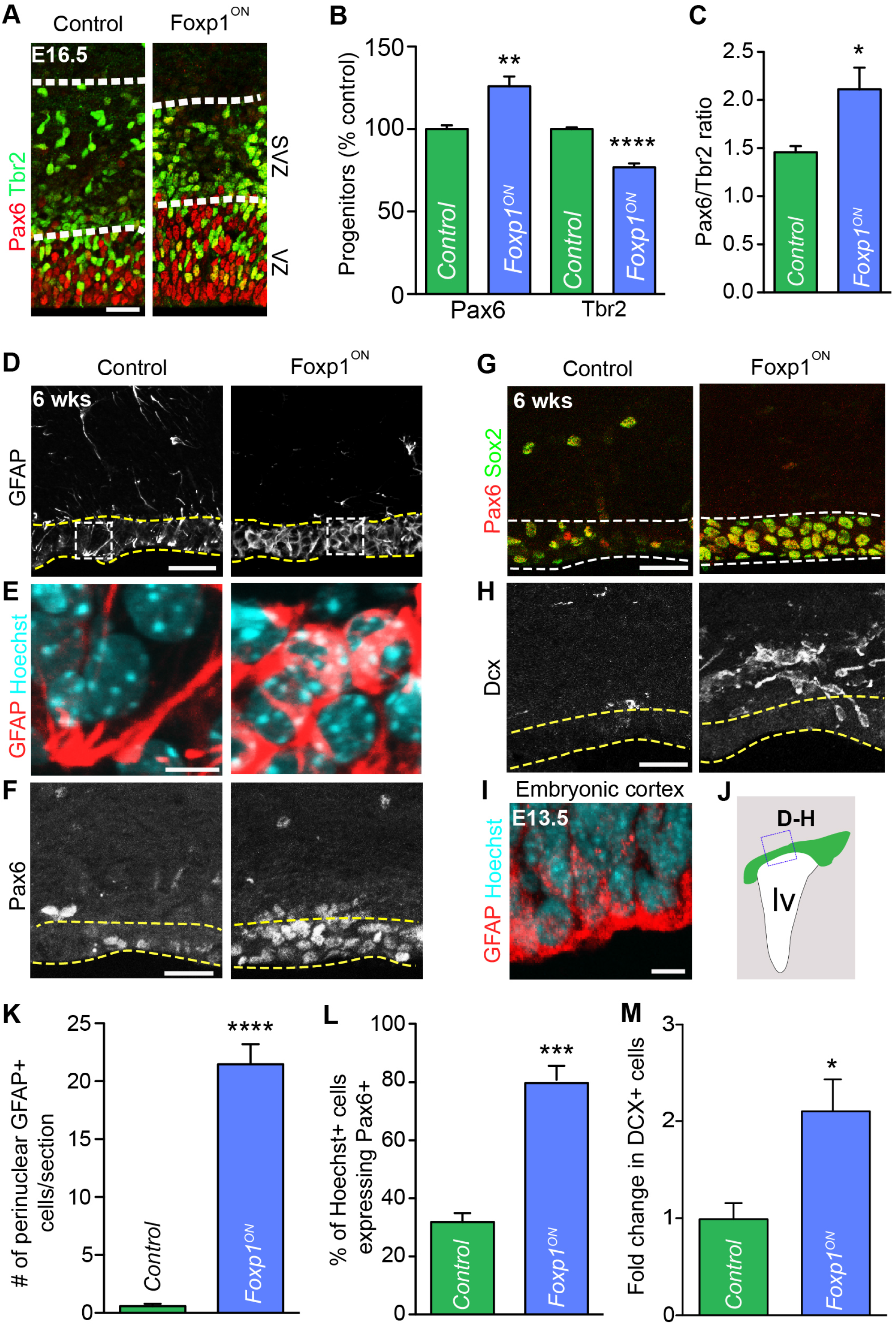
Foxp1 can sustain an aRG-like population into adulthood. (A) Immunostaining for Pax6 aRG and Tbr2 IP in E16.5 control and Foxp1^ON^ cortices. White dashed lines demarcate the approximate boundaries of the VZ and SVZ. (B-C) Relative number of Pax6+ and Tbr2+ cells in control and Foxp1^ON^ embryos at E16.5 and graphical representation of the Pax6 to Tbr2 ratio difference. (D-H) Immunostaining for GFAP, Pax6, Sox2, and Dcx in the dorsal lip of lateral ventricles in the forebrain of 6-week-old control and Foxp1^ON^ animals. Dashed yellow lines demarcate the approximate position of the SVZ region. Dashed white boxes in D denote regions displayed at higher magnification in E, which highlight the difference in GFAP localization in the SVZ region with nuclei labeled with Hoechst. (I) Comparative immunostaining for GFAP and Hoechst in wild type embryonic aRG at E13.5. Note the similarity in perinuclear GFAP staining seen in the postnatal Foxp1^ON^ forebrain. (J) Schematic depicting the region of the postnatal forebrain imaged in panels D-H. Green represents the region impacted by Emx1^Cre^-mediated recombination. (K) Number of cells with perinuclear GFAP staining per section within the lining of the lateral ventricle in 6-week control and Foxp1^ON^ animals. (L) Plot of the percentage of Hoechst cells lining the lateral ventricles that express Pax6. (L) Fold change in number of DCX+ cells in week 6 control and Foxp1^ON^ animals. All graphs display the mean ± SEM from multiple sections taken from 3-5 embryos or animals per condition. Significance was determined using the Student’s t-test. *p <0.05; **p <0.005; ***p <0.001; ****p <0.0001. VZ, ventricular zone; SVZ, subventricular zone; lv, lateral ventricle. Scale bars represent 50μm in panels A, D, F, G H and 5 μm in E, I. See also Figure S5.

Given the ability of Foxp1 to promote aRG maintenance in the late embryonic cortex, we next asked whether sustained expression of Foxp1 into adulthood could alter the transformation of these progenitors into adult neural stem cells. Emx1^Cre^-mediated recombination occurs in nearly all cells within the dorsal SVZ adjacent to the lateral ventricles (Young et al., 2007; Figure 5J), a major site of adult neurogenesis. At early postnatal stages, aRG are normally depleted as they give rise to astrocytes, ependymal cells, and adult neural stem cells of the SVZ (commonly referred to as B1 cells) (Lim and Alvarez-Buylla, 2016). B1 cells can be distinguished by their expression of Sox2 and low levels of Pax6, somatic location away from the edge of the ventricles, and thin GFAP+ processes protruding between ependymal cells to maintain contact with the ventricles (Figures 5D-5G). This profile is distinct from embryonic aRG which have high levels of both Sox2 and Pax6, tight somatic packing along the ventricles, and prominent perinuclear GFAP staining (Figure 5I and 5K). In 6-week-old Foxp1^ON^ mice, most GFAP+ cells lining the dorsal lateral ventricles appeared more similar to embryonic aRG than B1 cells (Figures 5D, 5E, 5I, and 5K). Moreover, the number of Pax6+ cells lining the ventricles significantly increased, from 32% of DAPI+ cells in controls to 80% in Foxp1^ON^ animals (Figures 5F, 5G, and 5L). The GFAP and Sox2/Pax6 costained cells also appeared to form a stratified neuroepithelium distinct from that seen in control animals (Figures 5D and 5G). Many of these Pax6+ cells expressed the ependymal cell marker Foxj1 (Figures S6A and S6B), suggesting either expanded production of these cells or possibly a conversion of some ependymal cells towards an aRG-like fate. Despite these changes, the ependymal lining of the ventricles, distinguished by S100 staining, was nevertheless preserved (Figure S6A).

Type B1 cells generate transit amplifying neuronal precursors (type C cells) which in turn produce neuroblasts (type A cells) and postmitotic neurons (Lim and Alvarez-Buylla, 2016). Type C cells, identified by their expression of the proneural protein Ascl1, were slightly reduced in Foxp1^ON^ animals (Figures S6A and S6B). At the same time, we found a 2-fold increase in the number of cells expressing Doublecortin (Dcx) (Figures 5H and 5M). While Dcx is frequently used as a marker of type A cells, it is also highly expressed by newborn neurons formed by embryonic aRG. We surmise that the increased numbers of Dcx+ cells seen here may thus reflect this mode of production. Collectively, these data show that Foxp1 misexpression alters the course of postnatal neurogenesis by maintaining a population of cells with aRG-like properties throughout embryogenesis and into adult life.

### Foxp1 is expressed by both aRG and bRG in the human fetal cortex

In light of the association of *FOXP1* mutations with human neurodevelopmental defects, we lastly investigated whether FOXP1 was similarly associated with neural progenitor maintenance and early, deep layer neurogenesis in the human cortex. FOXP1 was highly expressed within aRG as early as gestational week (GW) 13, and by GW14 also became prominent in newly-formed bRG located in the inner and outer subventricular zones (iSVZ and oSVZ; Figures 6A and 6B; arrowheads). The bRG identity of the FOXP1^+^ cells was confirmed by costaining for several molecular markers including SOX2, PAX6, HOPX, and SOX9 (Figures 6B and 6C). We further detected FOXP1 in mitotic PAX6+ pVIM+ bRG (Figures 6D and S7), a result that contrasted with the lack of overlap seen in mice (Figure 1G).

**Figure 6.**
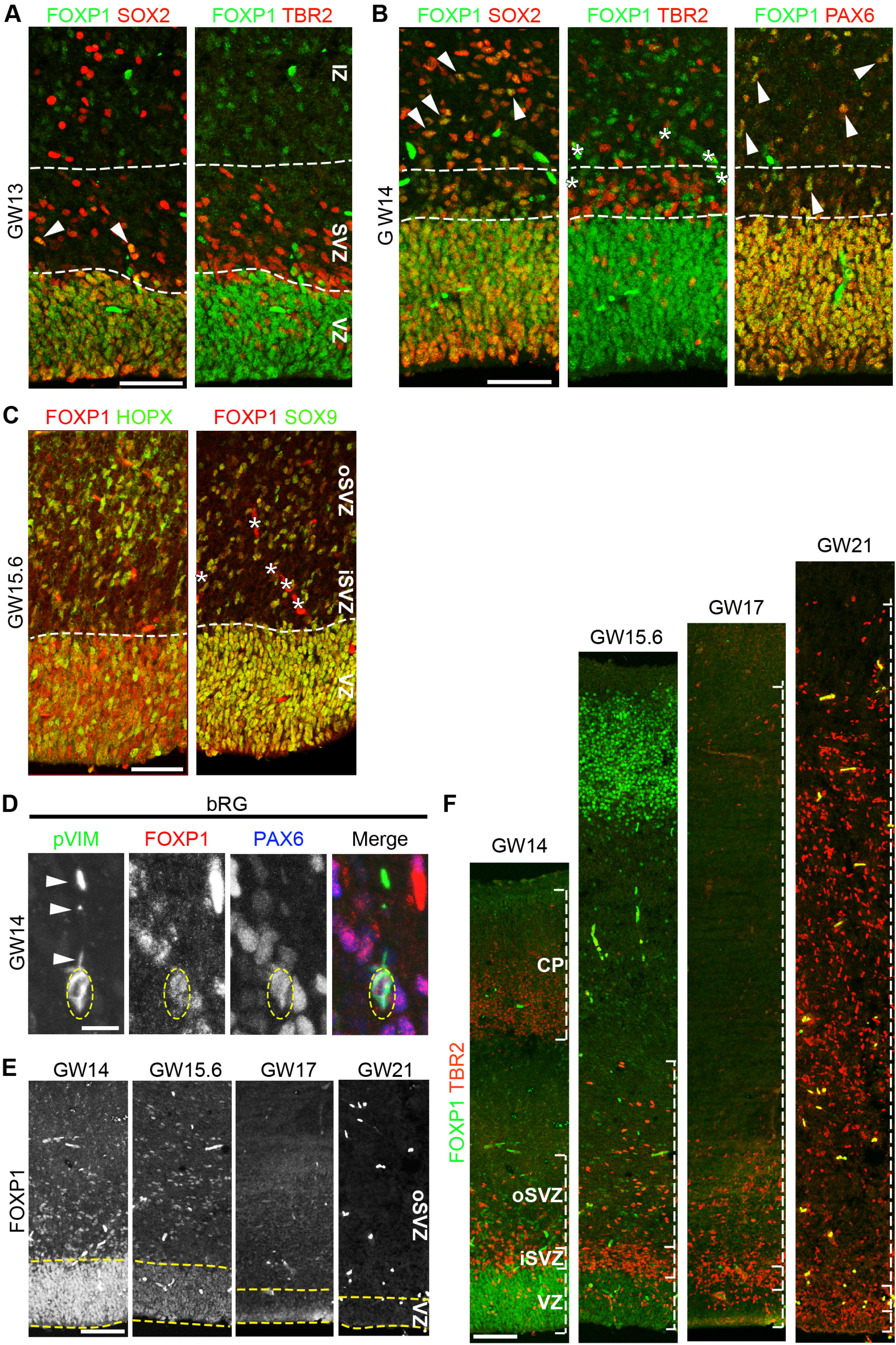
Foxp1 is expressed by both aRG and bRG in the human fetal cortex. (A) Immunostaining for FOXP1, SOX2, PAX6 and TBR2 in the VZ and SVZ at gestational weeks (GW) 13-14 in the human fetal cortex. Dashed white lines demarcate the boundaries of the VZ, SVZ and IZ. Arrowheads indicate FOXP1/SOX2 double positive cells. Asterisks indicate nonspecific staining in blood vessels (C) At GW15.6, nearly all FOXP1 cells in the SVZ express additional markers characteristic of bRG including HOPX and SOX9. (D) Immunostaining for FOXP1, phospho-VIMENTIN (pVIM) and PAX6 in the oSVZ at GW14. Dashed yellow oval denotes the nucleus in each channel. White arrows indicate pVIM staining in basal processes. (E-F) Analysis of FOXP1 levels from GW14-21 along with TBR2 immunostaining to demarcate the expansion in IP cells and the size of the iSVZ and oSVZ over time. VZ, ventricular zone; SVZ, subventricular zone; IZ, intermediate zone; iSVZ, inner subventricular zone; oSVZ, outer subventricular zone; bRG, basal radial glia; CP, cortical plate. Scale bars represent 50 μm in panels A-C, 10 μm in D, and 100 μm in E and F.

In mapping the distribution of FOXP1 over time in the human cortex (GW14-21), we found that FOXP1 levels progressively declined in both aRG and bRG between GW15-17, the time period over which the transition from deep to superficial layer neurogenesis is thought to occur (Nowakowski et al., 2016). At GW17, low levels of FOXP1 were nonetheless present in most, if not all bRG (Figure S7). However, FOXP1 expression in aRG and bRG was extinguished by GW21, coinciding with the start of gliogenesis (Figures 6E and 6F). Thus, as with mice, high FOXP1 levels demarcate the early period of neurogenesis in human cortical development, whereas lower levels are associated with late, superficial layer formation.

### Foxp1 misexpression generates ectopic bRG-like cells

Given the striking association of Foxp1 expression with bRG in the human cortex, we examined how its activity might relate to the formation of these cells using our mouse system. To distinguish the early effects of Foxp1 on aRG maintenance seen in our transgenic Foxp1^ON^ mice from later effects on bRG formation, we used *in utero* electroporation to deliver mouse Foxp1-IRES-nEGFP, human FOXP1-IRES-nEGFP, or nEGFP-only control expression plasmids into wild type mouse cortices at E13.5, the time at which endogenous bRG generation occurs (Wang et al., 2011). At 2 days post electroporation (E15.5), misexpression of either mouse or human Foxp1 more than doubled the fraction of GFP+ cells retained within the VZ compared to GFP-only controls, and reduced the number of GFP+ cells in the CP (Figures 7A and 7D). Both Foxp1 constructs also led to a striking appearance of Pax6+ cells throughout the IZ (Figure 7B). With mouse Foxp1, Pax6 was detected in ~85% of the GFP+ cells in the IZ, while human FOXP1 had a stronger effect with >95% of the GFP+ cells expressing Pax6 (Figures 7A-7C). Some GFP^+^ cells in the IZ also expressed neuronal markers such as Myt1l, indicating that at least some of the ectopic progenitors were capable of differentiating (Figure S7C).

**Figure 7.**
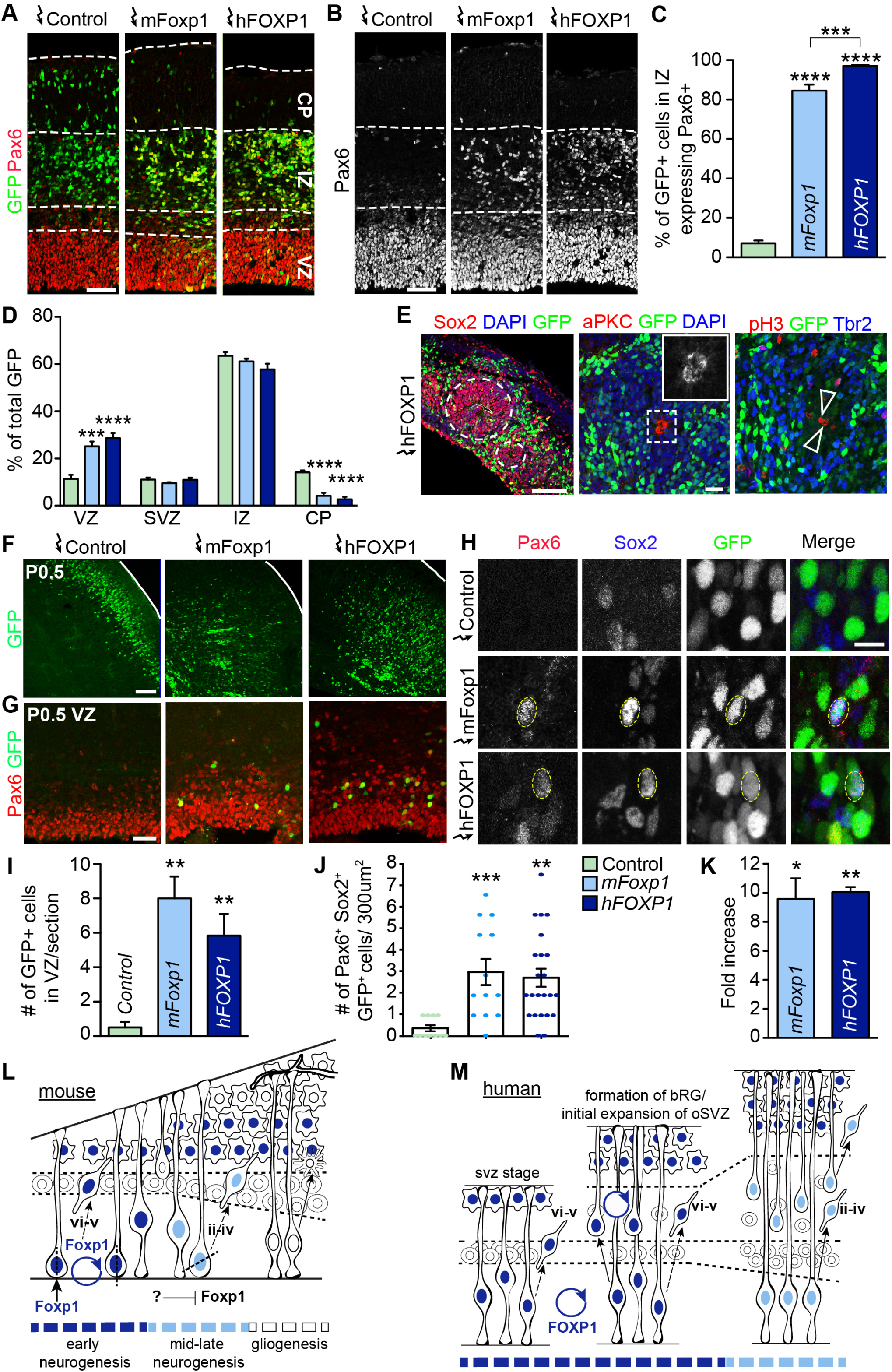
Foxp1 misexpression generates a population of bRG-like cells. (A-B) Immunostaining for GFP and Pax6 in cortices electroporated with control-IRES-GFP, mouse Foxp1-IRES-GFP (mFoxp1) and human FOXP1-IRES-GFP (hFOXP1) plasmids at E13.5 and collected 2 days post-electroporation (E15.5). White dashed lines demarcate boundaries of the VZ, SVZ, IZ and CP. (C) Quantification of the percentage of GFP+ cells in the IZ that express Pax6+ in control, mFoxp1 and hFOXP1-electroporated cortices. (D) Spatial distribution of GFP+ cells in cortices electroporated with indicated constructs at E13.5 and collected at E15.5. Plots in C and D display the mean ± SEM from multiple sections collected from 5-10 embryos per condition. (E) Immunostaining for Sox2, aPKC, phosphorylated Histone H3 (pH3) and Tbr2 in neural rosettes appearing in cortices 2 days after electroporation with human FOXP1. White dashed circles demarcate neural rosettes, solid line delineates a ventricle-like space. Dashed box and arrowheads highlight aPKC+ apical progenitor end feet and pH3+ mitotic cells along the inner lumen of the rosettes. (F) Distribution of GFP+ cells in cortices electroporated at E13.5 with control, mFoxp1, or hFOXP1 expression plasmids and collected at P0.5. (G) Immunostaining for Pax6 in the VZ of electroporated P0.5 cortices. (H) Analysis of ectopic Pax6/Sox2 double positive cells outside of the VZ found in P0.5 cortices electroporated with Foxp1 expression constructs. (I) Quantification of the number of GFP+ cells within the P0.5 VZ after electroporation with control, mFoxp1, or hFOXP1 plasmids. (J-K) Quantification of the number of Pax6/Sox2/GFP triple positive cells per 300 μm^2^ in Foxp-electroporated P0.5 cortices and resultant fold increases. Mean ± SEM from 4-12 sections/areas per condition are shown. (L) Model depicting the roles of Foxp1 regulating aRG maintenance and biasing neurogenesis towards early born fate in the mouse cortex. High levels of Foxp1 promote aRG identity (vertical, self-renewing divisions) and production of deep layer neurons. At mid-neurogenic stages, Foxp1 is downregulated in aRG (via an unknown mechanism). Subsequently, aRG divisions become more random and superficial layer neurons are generated. By the onset of gliogenesis, Foxp1 is no longer expressed by aRG. (M) Proposed model for the roles of FOXP1 during human corticogenesis. Initially, high levels of Foxp1 promote aRG identity and deep layer neurogenesis. By GW14, FOXP1 becomes associated with and may promote the formation and/or expansion of RG in the newly formed oSVZ. Over time, FOXP1 levels decline, permitting the onset of superficial layer neurogenesis and, ultimately, gliogenesis. FOXP1 decline may further influence bRG maintenance. Significance in panels D and I-L was determined using the Student’s t-test. ***p <0.001; ****p <0.0001. Scale bars represent 100 μm in panel E, 10 μm in H. VZ, ventricular zone; SVZ, subventricular zone; IZ, intermediate zone; CP, cortical plate. See also Figure S6.

In a third of all embryos electroporated with human FOXP1, we observed formation of neural rosette structures in the developing lateral cortex (Figure 7E). These rosettes exhibited apicobasal polarity characterized by apical aPKC antibody staining and formation of small ventricle-like spaces with pH3+ cells undergoing mitosis along the ventricular edge. Moreover, Tbr2+ IP were found around the circumference of these rosettes (Figure 7E). Only 1 out of 10 embryos electroporated with mouse Foxp1 and no EGFP vector control embryos displayed this phenotype. Together with the ectopic Pax6 results, these data illustrate differences in the progenitor-promoting activities of human vs. mouse Foxp1.

To assess the fate of the cells transfected with the different expression constructs, we analyzed the distribution of GFP+ cells in cortices one week post-electroporation, at P0.5. In control electroporations, the GFP+ cells were primarily located within the superficial layers of the CP with almost none found in the VZ (Figures 7F and 7G). However, after electroporation of mouse or human FOXP1, GFP+ cells became localized throughout the cortex, including a significant increase in GFP+ Pax6+ cells in the VZ (Figure 7G), again consistent with aRG retention within the postnatal VZ.

To determine whether Foxp1 misexpression also impacted bRG formation, we quantified coexpression of Pax6 and Sox2 within GFP+ cells located outside of the VZ. In control electroporations, we very rarely found GFP+ cells expressing Pax6 and Sox2 outside of the VZ (average 0.34 cells/300μm^2^; Figures 7H and 7I). However, following Foxp1 electroporation there were significant increases in GFP+ Pax6+ Sox2+ bRG-like cells, with a ~9-fold change seen with mouse Foxp1 and ~10-fold change with human FOXP1. Collectively, these data suggest that Foxp1 plays a critical role promoting aRG maintenance in both the mouse and human cortex, and may also contribute to formation and/or expansion of bRG.

## DISCUSSION

The progressive generation of neuronal subtypes in the cortex depends on both the maintenance of a pool of proliferating progenitors and their timely differentiation into first deep and then superficial layer neurons. Our studies identify Foxp1 as a critical regulator of these processes, acting to promote self-renewing vertical divisions that expand the aRG progenitor pool, while also favoring the generation of early-born deep layer neurons. Later stages in neurogenesis require Foxp1 downregulation, suggesting that its expression and activity helps define distinct windows of corticogenesis. The contributions of Foxp1 appear to be even more extensive in humans, as it is prominently associated with the early formation and proliferative expansion of bRG, raising the possibility that its progenitor maintenance functions may have been co-opted to facilitate the expansion of brain size across species. These observations may further explain how Foxp1 mutation in humans leads to a spectrum of neurodevelopmental defects.

### Foxp1 and the temporal competency model for corticogenesis

Neural progenitor transplantation studies, *in vitro* cell lineage analyses, and *in vivo* fate mapping experiments have all indicated that there is a window of plasticity during early neurogenesis in which aRG are competent to give rise to all cortical neuron subtypes (Desai and McConnell, 2000; Shen et al., 2006; Gao et al., 2014; Ma et al., 2017). This temporal competence model predicts that there must be factors expressed by early aRG that promote neural stem cell maintenance and bias the production of deep layer neurons. As neurogenesis proceeds, the actions of these factors would need to be reduced or counterbalanced by opposing factors to enact a switch to superficial layer neurogenesis. Indeed, several factors have been implicated in this switching process including for early fates: Otx1, Foxg1, CoupTF1I/II, Fezf2 and Ikaros (Frantz and McConnell, 1996; Okano and Temple, 2009; Tuoc et al., 2009; Alsio et al., 2013; Guo et al., 2013; Toma and Hanashima, 2015), and Brn1/2 and Cux1/2 for later fates (Franco et al., 2012; Dominguez et al., 2013). However, most of these factors appear to exhibit some but not all of the cardinal features of a temporal determinant, namely restricted expression in progenitors during defined time windows of neurogenesis, a requirement for that phase of neurogenesis, and a capacity to alter the intervals of neuron production when misexpressed. In this regard, Foxp1 stands out as an intriguing candidate as its expression demarcates the period of deep layer neuron production, and its function is both necessary and sufficient for this process.

Our electroporation experiments show that acute misexpression of Foxp1 in aRG during the time window of superficial layer neurogenesis can elicit the generation of the earliest born Layer I cells, as well as deep layer neurons. Similar results were observed in transgenic Foxp1^ON^ mice, where deep layer neurons continued to be generated at later stages of embryogenesis and early postnatal life. However, despite the extended production of deep layer cells, superficial layer neurogenesis nevertheless continued in the face of ectopic Foxp1 expression. These findings suggest that whilst Foxp1 can broaden the neurogenic competence of aRG, its functions alone do not prohibit formation of later born cell types. Supporting this conclusion, forced expression of Foxp1 in postmitotic neurons using Nex1^Cre^-mediated recombination neither stimulated the formation of deep layer neurons nor blocked superficial fates.

### Potential downstream targets of Foxp1 in corticogenesis

Foxp1 exhibits diverse activities in the developing nervous system, reflecting the different cell types in which it is expressed and phases of differentiation with which it is associated. Moreover, Foxp1 is known to exist in multiple splice isoforms, and act in both homodimeric and heterodimeric repressor complexes in conjunction with the related proteins Foxp2 and Foxp4 (Wang et al., 2003; Li et al., 2004). The downstream targets of Foxp1 remain unclear though progress towards this goal is starting to be made considering its postmitotic context. In postnatal cortical neurons, Foxp1 can repress genes involved in neurogenesis, neuronal migration, and synaptogenesis (Usui et al., 2017). Similarly, studies using cultured late stage cortical progenitors and differentiated neurons have proposed that Foxp1 directly represses the Notch receptor ligand Jagged1 to promote neuronal differentiation (Braccioli et al., 2017).

It is unclear how these findings relate to the actions of Foxp1 in early neural progenitors which we have focused on in our studies. The ability of Foxp1 to promote aRG maintenance suggests that some of its key targets likely include factors that stimulate cell differentiation as well as those that control the symmetry of progenitor cell divisions. Vertical divisions are tightly regulated by the spindle orientation machinery present in early progenitors. As neurogenesis proceeds, this regulation is relaxed and divisions become more random (Paridaen and Huttner, 2014), coincident with Foxp1 downregulation. Foxp1 could potentially be involved in the repression of factors such as Inscutable that can disrupt the mitotic spindle and thereby randomize the angle of progenitor divisions (Lancaster and Knoblich, 2012). Likewise, the ability of Foxp1 to promote deep layer neuronal character may result from its ability to repress factors associated with superficial neuronal fates.

One factor that was consistently impacted by Foxp1 gain and loss was Pax6, one of the earliest markers of cortical progenitors. Pax6 expression is maintained in aRG throughout embryonic neurogenesis, and is also utilized by neural stem and progenitor cells in the SVZ of the postnatal and adult forebrain. In humans and non-human primates, Pax6 is further associated with the formation and maintenance of bRG cells. At each stage of development, Foxp1 misexpression substantially increased Pax6 levels suggesting that many of its progenitor-promoting activities could be attributed to this induction. There are striking similarities in the consequences of *Pax6* and *Foxp1* deletions, as mice lacking either gene exhibit more oblique/horizontal divisions, reduced aRG proliferation, and, at later stages, decreased neuron production (Ypsilanti and Rubenstein, 2016). Likewise, ectopic Pax6 expression can impede neuronal differentiation and induce the formation of bRG-like cells in the mouse cortex much like Foxp1 (Wong et al., 2015).

### Contributions of Foxp1 to human cortical development

Significant progress has been made comparing the transcriptomic landscapes of human and mouse cortical progenitors (Florio et al., 2015; Johnson et al., 2015; Pollen et al., 2015). These analyses have identified novel bRG markers and provided insights into how human bRG are regulated and what distinguishes them from other progenitor populations. However, to date, few factors have been identified that differentiate between mouse and human bRG. Our studies show that Foxp1 is not expressed in mouse bRG but is prominently expressed in human bRG, particularly during the early phase of corticogenesis. Comparisons of mouse and human bRG and have demonstrated that mouse bRG are less proliferative, have a reduced capacity for self-renewal, and are transcriptionally comparable to IP whereas human bRG most closely resemble aRG (Johnson et al., 2015).

Little is currently known about the mechanisms that regulate bRG formation and expansion to create the enlarged oSVZ progenitor compartment seen in the human and non-human primate brain. Our experiments suggest that Foxp1 contributes to these processes. When ectopically expressed in the mouse cortex at mid-neurogenesis, Foxp1 elicited the formation of abventricular Pax6+ cells, many of which gave rise to deep layer neurons. Nevertheless, a subset of the Foxp1-transfected cells located outside of the VZ maintained both Pax6 and Sox2 expression, a characteristic feature of bRG, for at least a week after electroporation. These results suggest that Foxp1 may also enable long-term maintenance and expansion of bRG over time. This function fits with our findings that Foxp1 is present in both aRG and bRG in the human cortex during the peak period of oSVZ expansion between GW14 and GW17 (Nowakowski et al., 2016).

It is notable that the high to low gradation in Foxp1 that we have observed in mouse and human aRG across time also holds true for bRG. This differential expression could impact both the proliferation and self-renewal capacity of the bRG at different stages of development, while also influencing their capacity to give rise to deep vs. superficial layer neurons.

While increasing Foxp1 expression promoted both aRG maintenance and bRG expansion in mice, we did not observe dramatic increases in brain size in any of our experiments. Some of this lack of effect may reflect technical details in our experiments, for example the early timing at which Foxp1 is first expressed in the Foxp1^ON^ mice favoring aRG maintenance vs. bRG production, as well as mosaicism and transience of the effects achieved with DNA electroporation. However, it seems very likely that whilst Foxp1 can increase bRG numbers, collaborating factors are required to significantly influence brain growth. The oSVZ is a large compartment composed not only of bRG, but also IP (whose formation is blocked by sustained Foxp1 expression) and other cells that together create a microenvironment permissive for proliferation and expansion, particularly in the human cortex (Lui et al., 2014).

### Pathological implications of Foxp1 dysregulation

Foxp genes are members of the evolutionarily ancient Fox family, and their activities have been linked to the acquisition of human specific traits such as language, speech and intellectual capabilities (Hannenhalli and Kaestner, 2009). However, a consequence of the selective pressures that permitted language and cognition in humans also made us vulnerable to cognitive disorders such as intellectual disability and ASD (Lepp et al., 2013). Copy number variants, haploinsufficiencies and de novo point mutations in *FOXP1* have been found in patients exhibiting ASD and speech impairment (Hamdan et al., 2010; Horn et al., 2010; O’Roak et al., 2011; Le Fevre et al., 2013; Meerschaut et al., 2017). Intriguingly, increased expression of *FOXP1* has also been reported in patients with ASD (Chien et al., 2013).

Mouse models have linked Foxp1 in striatal neurons with ASD-like behavior and transcriptional regulation of autism-related pathways (Araujo et al., 2015; Bacon et al., 2015). Next generation sequencing studies further show that many genes implicated in ASD are highly coexpressed during human cortical development and can be grouped into modules that involve distinct biological functions including early transcriptional regulation (Parikshak et al., 2015). These studies underscore the importance of understanding the spatial and temporal context of specific ASD susceptibility genes in order to ascertain their role in brain development. Much of the research involving Foxp1 and human neurodevelopmental disorders has thus far focused on the functions of Foxp1 in postmitotic neurons. Our findings that experimentally induced changes in Foxp1 levels can alter the maintenance of neural progenitors, production of bRG, and time windows of deep vs. superficial layer neurogenesis add new perspectives in how Foxp1 mutations could impact the growth, cytoarchitecture, and evolution of the human cerebral cortex.

## ACKNOWLEDGEMENTS

We are grateful to K. Adams, H. Martin, E. Morrisey, D. Rousso, B. Van Handel, M. Mall and M. Wernig for materials; K. Phan and N. Vishlaghi for technical assistance; the UCLA Broad Stem Cell Research Center for microscopy and other resources; K. Adams, S. Butler, L. De La Torre-Ubieta, D. Geschwind, M. Harrison, M. Watanabe, and members of the Novitch and Butler laboratories for invaluable discussions and comments on the manuscript. This work was supported by awards from the UCLA Broad Stem Cell Research Center, and grants to B.G.N. from the NIH (NS089817, 1R01NS085227, and R01NS072804) and the California Institute for Regenerative Medicine (DISC1-08819), and to H.O.T from the NIH (AI059447), CPRIT (RP100612, RP120348) and the Marie Betzner Morrow Centennial Endowment. C.A.P. was supported by the UCLA-California Institute for Regenerative Medicine Training Grant (TG2-01169), and J.D.D was supported by the Lymphoma Research Foundation.

## AUTHOR CONTRIBUTIONS

Conceptualization, CAP and BN.; Methodology, CAP and BN.; Investigation, CAP, DM, and AM.; Resources, HT, JD and HH.; Writing – Original draft, CAP and BN.; Writing – Review and Editing, CAP and BN.; Visualization, CAP.; Supervision, BN.; Project administration, CAP.; Funding acquisition, BN.

## STAR METHODS

Detailed methods are provided and include the following:

- Key resources table.
- Contact for reagent and resource sharing.
- Experimental Model and subject details.

- Animal preparation and tissue analysis.
- Human fetal tissue.
- Method details.

- BrdU incorporation analyses.
- Plasmid expression and constructs.
- In utero electroporation.
- Microscope imaging.
- Quantification and Statistical analysis.

- Cell and protein staining quantification.
- Statistical analysis
- Data software and availability.

**Table 1.**
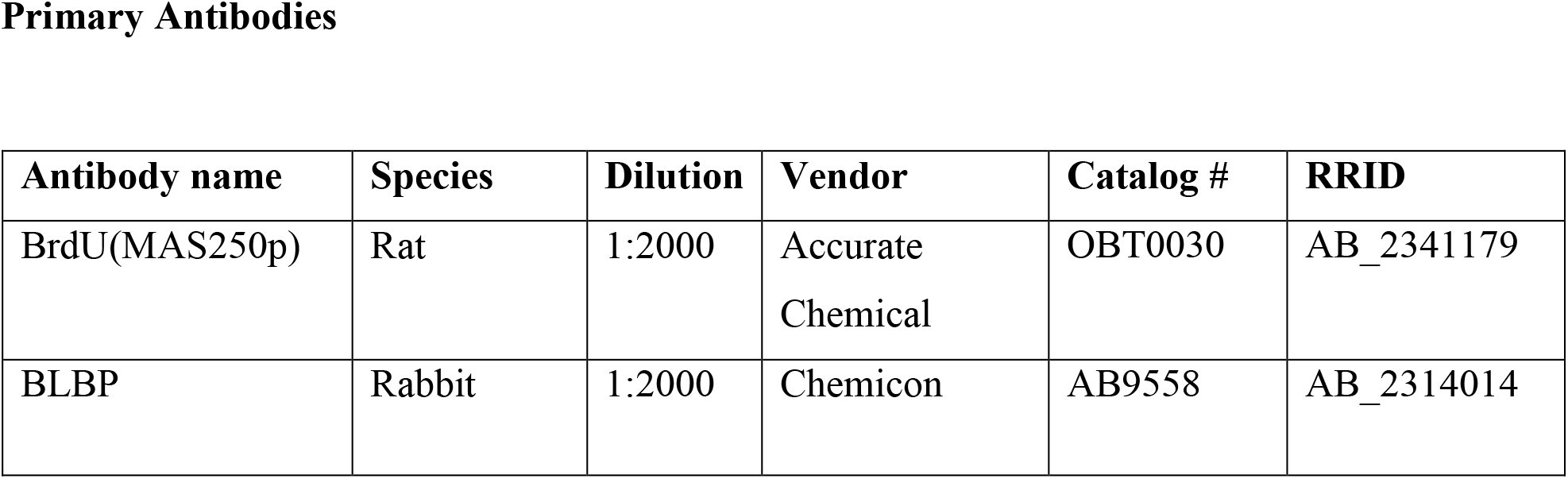
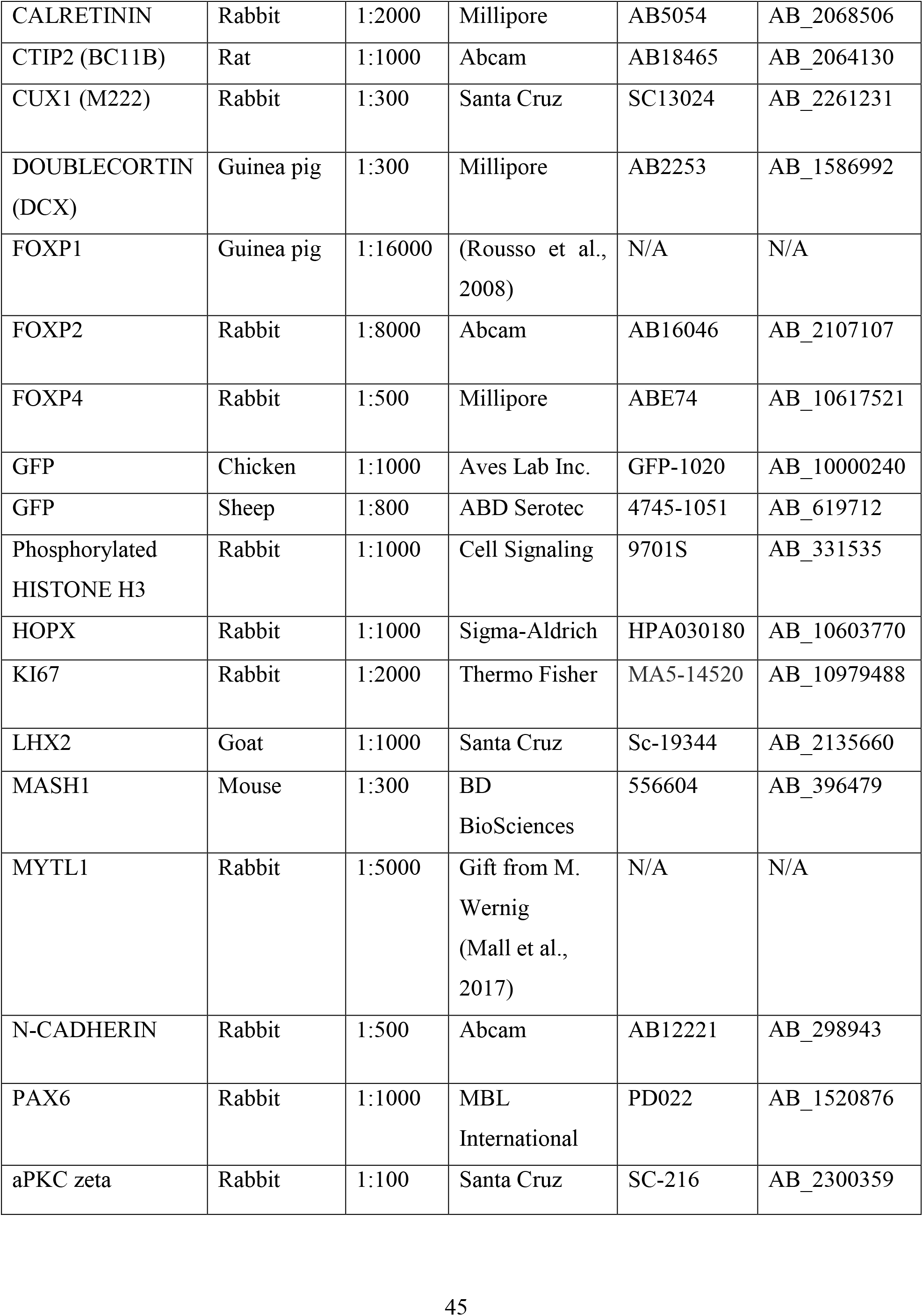
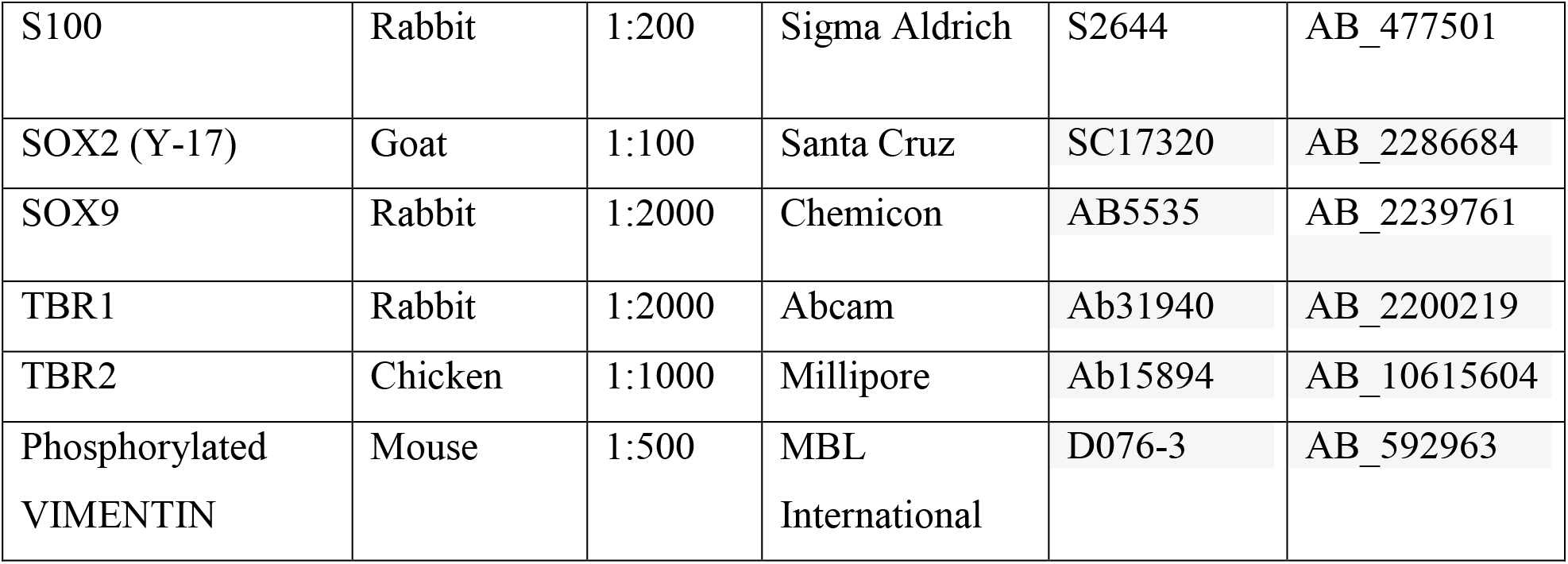
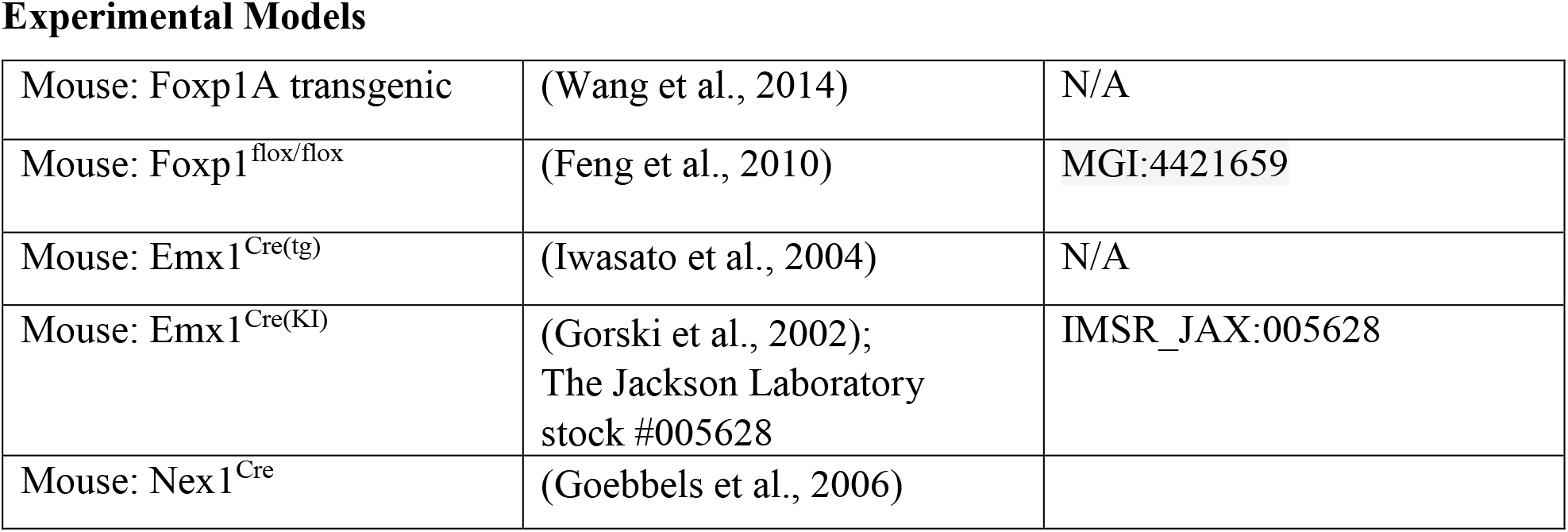
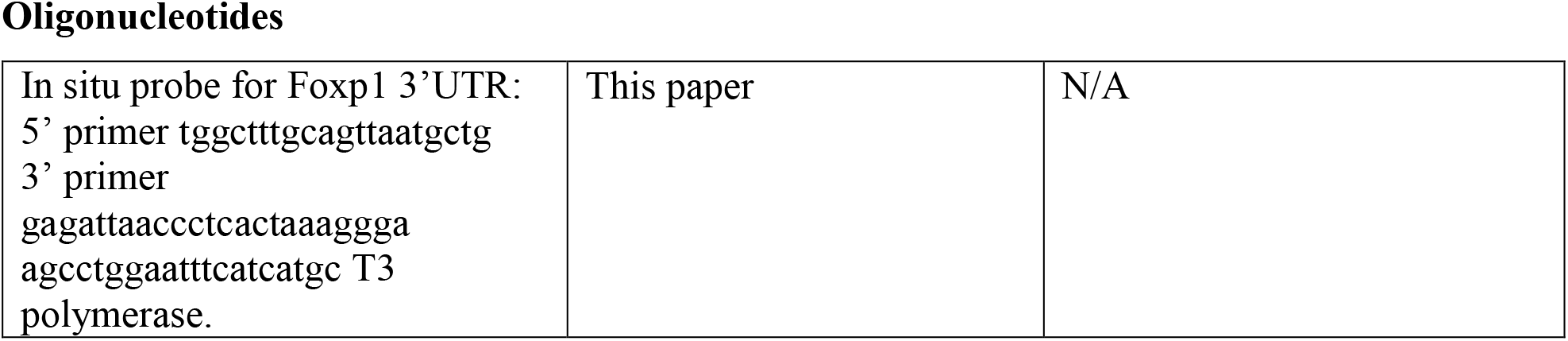
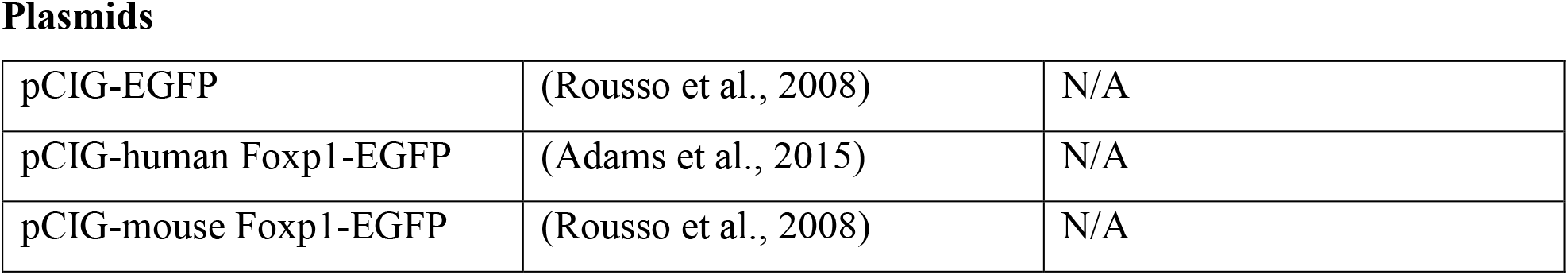
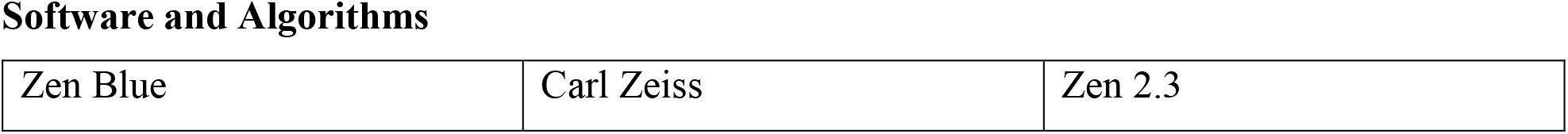
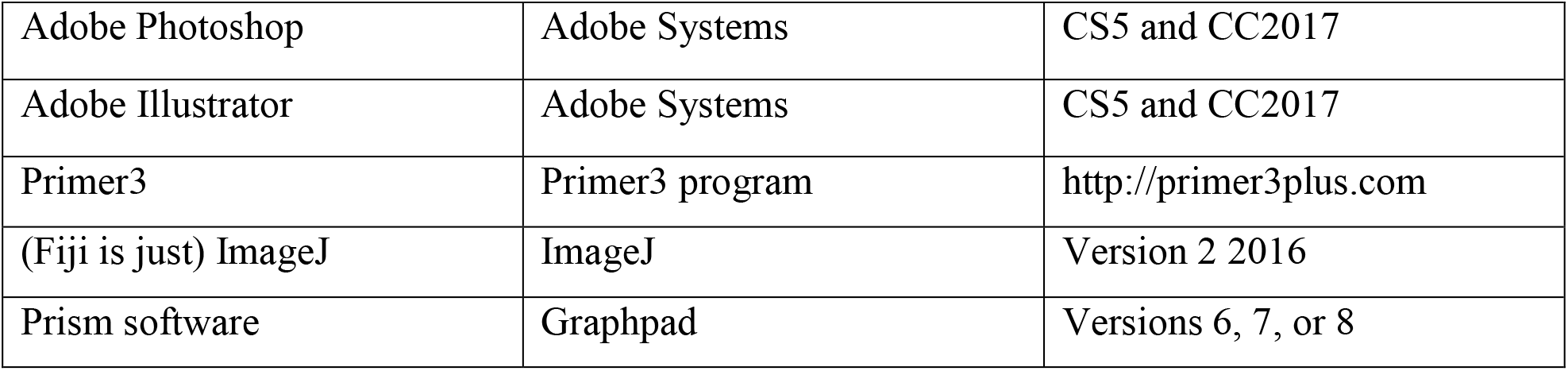
KEY RESOURCES TABLE

## CONTACT FOR REAGENT AND RESOURCE SHARING

Additional information and requests for resources and reagents should be directed to the Lead Contact, Bennett Novitch.

## EXPERIMENTAL MODEL AND SUBJECT DETAILS

### Animal preparation and tissue analysis

*Foxp1^tg/tg^*, *Foxp1^fl/fl^*, *Foxp1*^*fl*/+;^ *Emx1*^*Cre/*+^ mice were maintained as previously described (Gorski et al., 2002; Iwasato et al., 2004; Goebbels et al., 2006; Feng et al., 2010; Wang et al., 2014) following UCLA Chancellor’s Animal Research Committee husbandry guidelines. Embryonic cortices were fixed in 4% paraformaldehyde overnight. Postnatal and adult animals were perfused according to UCLA guidelines, and brains were fixed overnight. Tissues were cryosectioned and processed for immunohistochemistry or in situ hybridization as previously described (Pearson et al., 2011; Rousso et al., 2012). Primary antibodies are listed in the Key Resources table. A probe against the 5’UTR of mouse *Foxp1* was generated using PCR.

### Human fetal tissue

Experiments were performed with prior approval from the research ethics committees at the UCLA Office of the Human Research Protection Program, the University of Tübingen (institutional review board [IRB]#323/2017BÜ2), and Novogenix Laboratories. Embryonic tissues were obtained with informed consent as discarded materials resulting from elective legal terminations. Samples were de-identified in accordance with institutional guidelines. Specimen ages for this study are denoted as gestational weeks as determined by the date of the last menstrual period or ultrasound and confirmed by analysis of developmental characteristics. Samples were processed as described above.

## METHODS DETAILS

### Plasmid expression constructs

Generation of mouse and human Foxp1 expression vectors was conducted as previously described (Rousso et al., 2008; Adams et al., 2015). In brief, the coding region of each gene was amplified by PCR and cloned into a Gateway compatible version of the pCIG expression vector, which carries an IRES-nuclear-EGFP reporter.

### In utero electroporation

Electroporations were performed as previously described (Cruz-Martin et al., 2010). A solution containing 1.0μg/μL of plasmid DNA (pCIG or pCIG-Foxp1) and 0.05% Fast Green was injected in the left telencephalic vesicle at E13.5, and 5 square electric pulses (40V, 50 ms long) were delivered at 500 ms intervals using a BTX Harvard Instruments electroporator. Brains were recovered at E15.5 or at birth (P0.5), fixed in 4% paraformaldehyde, cryosectioned, and used for immunohistochemical staining analysis.

### Microscope Imaging

Confocal images were acquired using Zeiss LSM 780 or LSM 800 confocal microscopes and Zen blue or black software. DIC imaging of in situ hybridizations were collected using a Zeiss Axioimager microscope and Axiovision software. Images were processed and compiled using Adobe Photoshop with image adjustments applied to the entire image and restricted to brightness, contrast and levels. Images shown in figures as comparisons, e.g. intensity levels, were obtained and processed in parallel using identical settings. Composite images were assembled using Adobe Illustrator software.

## QUANTIFICATION AND STATISTICAL ANALYSES

### Cell and protein staining quantification

For each experiment, the number of labeled lateral cortex cells within a ~200 μm x radial width area per section was quantified from at least three 16 μm sections sampled at ~100 μm intervals along the rostrocaudal axis. The number of cells in each Foxp1 condition was normalized to littermate control embryos. For electroporation experiments, GFP+ cells were counted and the percentage of those cells expressing specific markers calculated. For BrdU birthdating experiments, the total number of BrdU^+^ cells within either Layers V/VI or II/III-IV were counted and the percentage of BrdU+/Tbr1+ or Lhx2+ cells calculated. For the later embryonic incorporation experiments, the total number of BrdU+ cells in the CP were counted and the number of double positive cells quantified. For mitotic division angle analyses, high magnification images of dividing cells in anaphase along the ventricular wall were taken and angle was measured relative to apical surface. Between 80-100 cells per condition were measured. Intensity values relative to background staining were measured using Fiji software, using images from each condition that were measured using identical settings.

### Statistical Analyses

The normality of each data set was determined using Graphpad Prism software and the appropriate parametric (normal distribution) or non-parametric (uneven distribution) test was applied as indicated in the Figure legends. Unpaired t-tests and Mann-Whitney tests were calculated using Prism software. Wilcoxon signed rank test was applied in the case where control values were identical (Figure S5B). Significance was assumed when p < 0.05. Throughout the manuscript, the results of statistical tests (p values and n numbers) are reported in the figure legends. Signifiers used are as follows: p > 0.05, ns; * p > 0.05; ** p > 0.01; *** p > 0.001; **** p > 0.0001. Unless otherwise indicated, all data are presented as mean± SEM.

## DATA AND SOFTWARE AVAILABILITY

All statistics and graphs were generated using Graphpad Prism 6, 7, or 8 software. See Key resources table for information regarding other software used.

